# Exploiting regulatory heterogeneity to systematically identify enhancers with high accuracy

**DOI:** 10.1101/250241

**Authors:** Hamutal Arbel, William W. Fisher, Ann S. Hammonds, Kenneth H. Wan, Soo Park, Richard Weiszmann, Soile Keränen, Clara Henriquez, Omid Shams Solari, Peter Bickel, Mark D. Biggin, Susan E. Celniker, James B. Brown

## Abstract

Identifying functional enhancers elements in metazoan systems is a major challenge. For example, large-scale validation of enhancers predicted by ENCODE reveal false positive rates of at least 70%. Here we use the pregrastrula patterning network of *Drosophila melanogaster* to demonstrate that loss in accuracy in held out data results from heterogeneity of functional signatures in enhancer elements. We show that two classes of enhancer are active during early *Drosophila* embryogenesis and that by focusing on a single, relatively homogeneous class of elements, over 98% prediction accuracy can be achieved in a balanced, completely held-out test set. The class of well predicted elements is composed predominantly of enhancers driving multi-stage, segmentation patterns, which we designate segmentation driving enhancers (SDE). Prediction is driven by the DNA occupancy of early developmental transcription factors, with almost no additional power derived from histone modifications. We further show that improved accuracy is not a property of a particular prediction method: after conditioning on the SDE set, naïve Bayes and logistic regression perform as well as more sophisticated tools. Applying this method to a genome-wide scan, we predict 1,640 SDEs that cover 1.6% of the genome, 916 of which are novel. An analysis of 32 novel SDEs using wholemount embryonic imaging of stably integrated reporter constructs chosen throughout our prediction rank-list showed >90% drove expression patterns. We achieved 86.7% precision on a genome-wide scan, with an estimated recall of at least 98%, indicating high accuracy and completeness in annotating this class of functional elements.

**Significance Statement:** We demonstrate a high accuracy method for predicting enhancers genome wide with > 85% precision as validated by transgenic reporter assays in *Drosophila* embryos. This is the first time such accuracy has been achieved in a metazoan system, allowing us to predict with high-confidence 1640 enhancers, 916 of which are novel. The predicted enhancers are demarcated by heterogeneous collections of epigenetic marks; many strong enhancers are free from classical indicators of activity, including H3K27ac, but are bound by key transcription factors. H3K27ac, often used as a one-dimensional predictor of enhancer activity, is an uninformative parameter in our data.

## Introduction

Enhancers are ∼100 – 1,000 bp *cis*-regulatory elements that direct spatial and temporal patterns transcription in metazoans. Definitive epigenetic signatures of enhancer elements have been challenging to identify. A number of computational tools have been developed to predict enhancer elements from chromatin state and transcription factor *in vivo* DNA binding information (1-12). Tools that attempt to measure predictive accuracy using only indirect evidence of enhancer activity, e.g. enrichment in H3K27ac or p300, often display excellent accuracy by these limited criteria (1, 3, 13, 14). When algorithems are benchmarked on held-out *in vivo* tests of functional enhancer activity, however, positive predictive power on genome-wide scans in metazoan systems has been lower than expected. In most cases, precision rarely exceeds 40% (13-15). By targeting transcription factors in a specific biological proceces, however, a higher precision of 56% was achived in a randomly selected sample through transiant transfection (16). Higher precision has also been reported when tests were confined to the top of prediction rank list (17), but such numbers are unlikely to represent the precision of the prediction set as a whole.

There are several possible explanations for the poor accuracy of current enhancer prediction algorithms. The transient *in vivo* enhancer assays often employed to test predictions may suffer a high false-negative rate due to the loss of local chromatin context. Alternatively, the data provided to the prediction algorithms might be insufficient. Features such as H3K27ac and p300 can partially distinguish active enhancers (18, 19), but it remains unclear whether any chromatin mark or combination of chromatin marks and p300 uniquely identifies enhancers among all sequences in a genome (16, 20). Indeed, enhancers that lack H3K27ac and admit patterns of hyper methylation are essential during early vertebrate development (21). Hence, there may be more than a single class of genomic element that drives patterned expression, or, more precisely, the term “enhancer” may encapsulate a mechanistically diverse class of functional elements. Transcription factor (TF) occupancy is a better predictor of enhancer activity than canonical chromatin marks (including H3K27ac, H3K4me1, and H3K4me3) in mouse and humans (16). Thus mechanistic sub-types of functional enhancer elements may emerge from distinct patterns of TF occupancy and chromatin context.

To test the possibility that heterogeneity among enhancers is a major reason for the difficulty in predicting enhancers, we have exploited the pregrastrula *Drosophila* embryo network. A cohort of ∼30 spatially patterned TFs drive body patterning in concert with another 30 or so ubiquitously expressed sequence specific TFs (22-32). Embryo symmetry is broken along the anterior-posterior (A-P) axis and dorsal-ventral (D-V) axis by two separate sets of maternally deposited TFs. Over a 90 minute period corresponding to developmental stages 4 and 5, these proteins act in concert with zygotically expressed A-P and D-V TFs to refine initially broad patterns of transcription into narrower striped patterns that define the basic segmental body plan of the fruit fly (33). The pregrastrula fly network is thus a particularly well defined model system for studying the relationship between TF DNA binding and spatially patterned enhancer activity.

We have tested the utility of a wide range of data for predicting enhancers, including *in vivo* DNA binding patterns for 22 pregrastrula TFs, a variety of chromatin marks, evolutionary conservation, whole embryo mRNA-seq, and RNA polymerase II location. Using a test set of nearly 8,000 genomic regions whose enhancer activity had been determined in transgenic assays in whole-embryos (34, 35), we applied supervised machine learning to identify enhancer sequences active in pregrastrula embryos. Verified enhancers were separated into two approximately equally sized groups based on the reproducibility with which they were correctly predicted in multiple runs of a random forest. A model trained using the set of enhancers that were reproducibly classified correctly has >98% predictive accuracy when tested on a balanced set of known enhancer positive and negative genomic regions. In contrast, the other set of training enhancers generated models that predicted no better than random. Subsequent analyses revealed that the well predicted class of enhancers are near genes that show a strong tendency to be involved in controlling segmentation and other developmental processes and to be expressed in many cells of the embryo. The poorly predicted enhancers are without obvious ties to the control of segmentation and tend to be expressed in less than 15% of cells. By focusing on the well predicted class of enhancers, which we term segmentation driving enhancers (SDEs), we find that TF DNA binding is highly predictive, whereas histone modifications and the remaining features tested have little or no additional predictive power.

In a *de novo*, genome wide prediction, we predict approximately 1,640 SDEs in the early embryo that cover 1.6% of the euchromatic genome. As validated by an *in vivo* transgenic reporter gene assay, this set are predicted with 98% estimated recall and 95% precision. Unlike most previous studies, we concentrated validation away from the top of our rank list to increase the likelihood of identifying false positives and to improve our power to compute accurate error rates. Importantly, we show that our model performance is driven by the need to treat SDEs separately from other enhancer elements, rather than the properties of a specific computational method—naïve Bayes and logistic regression perform as well as more complex models after conditioning on the SDE set. This demonstrates the prediction of a specific class of enhancers with sufficient precision to enable their identification genome-wide.

## Results

### Data, Feature and feature selection

Transgenic reporter data for enhancer activity in *Drosophila* embryos were combined from two sources. Kvon *et al.* (34) had conducted a semi-automated screen of the reporter gene expression patterns driven by 7705 genomic regions (http://enhancers.starklab.org) at multiple stages throughout embryogenesis. While this high throughput assay allowed an unprecedented number of genomic areas to be tested, the small number of embryos per collection plate led to increased misclassifications in the data. The activity of an additional 282 genomic segments was determined by the Berkeley Drosophila Transcription Network Project (BDTNP) (35) (unpublished data). Altogether, 7987 genomic regions were examined and 731 were experimentally found to drive reporter gene expression in *Drosophila* embryonic stages 4-6 (36) (**SI table 1**). By manually comparing the activity of overlapping genomic regions in the BDTNP database with the larger data from Kvon et al. we estimate a 10% false negative rate in the latter.

Features used in the initial model included ChIP-chip data for 20 of the TFs that pattern transcription along the A-P and D-V axes of the embryo (37-39); ChIP-seq data for the ubiquitous TFs ZLD and Zeste; 45 chromatin proteins and histone modifications (40); DNase accessibility data (41-43); and evolutionary conservation scores (44-46). Also considered were the presence of bidirectional RNA transcripts; exon and intron coverage; distance to RNA Polymerase II ChIP-chip binding peaks; and distance to transcription start sites. A summarized list of features is presented in **Table 1**. For a full list and description please refer to **SI table 2** and **methods.**

**Table 1:** A summarized list of features used in initial model

| Category | Features included |
| --- | --- |
| Histone and Histone modifications | *H3, H3K18ac, H3K27ac, H3K27me3, H3K36me3, H3K4me1, H3K4me3, H3K9ac, H4K5ac, H4K8ac* |
| A-P regulatory transcription factors | *bcd, cad, gt, hb, kni, kr, hkb, tll, d, ftz, prd, run, slp* |
| D-V regulatory transcription factors | *da, dl, mad, med shn, sna, twi* |
| Ubiquitous Transcription Factor | *z, zld, sum of all transcription factors* |
| DNA data | Conservation, DNA accesibilty, Distance to Pol. II, Distance to TSS, bidirectional-RNA transcription |
| Exon/intron data | Exons, Coding exons and introns coverage/presence |

With this data we trained and tested Random Forest, a supervised machine learning approach based on an ensemble of decision trees (47-49). To reduce parameter number and prevent overfitting, we culled input features **(methods)**. We found that TFs and histone modification data were sufficient to minimize the error rate. We note that while DNase accessibility did not contribute to Random Forest predictive power in the presence of TF binding data, it significantly improved predictive power in its absence, suggesting multi-collinearity in the data. Conversely, conservation scores did not contribute to the predictive power in any fitted model, and the error rate utilizing solely conservation scores was ∼50%, suggesting that conservation is not a feature of enhancers in the *Drosophila* embryo.

### Heterogeneity among enhancer elements

With our optimal feature set, our error rate in a single forest as defined by misclassification was nearly 30%. The performance of the Forest voting probabilities as indicated by the area under the ROC curve, AUC=0.82 (**Fig. 1A**), is very similar to that in previously published work (16, 17), implying a similarly modest success rate. However, while this overall predictive power falls short of that required for predicting enhancers genome wide, we noticed that some enhancers were consistently correctly classified, while others were consistently misclassified. Hypothesizing that the model’s poor performance may be due to heterogeneity in the enhancer set, enhancers were separated into two classes. Class I contained the 358 enhancer segments that were correctly classified at least 75% of the time and class II containing the 373 that were not. When class II enhancers were excluded from the test sample, the single forest error rate drops to ∼3%, and the area under the ROC curve is ∼0.99 (**Fig. 1A**). When Class I enhancers were excluded from the test sample instead, errors of a single forest are ∼40%, and the ROC curve indicated performance only marginally better than random guessing (**Fig. 1A**). To establish that the enhancer heterogeneity is data-driven and not an artifact of our choice of method, logistic regression and naïve bays models of the data were also constructed. In both cases the removal of the class II enhancer set significantly improves the model’s predictive power (**Fig. 1A**). Interestingly, the effect of retaining and removing class I and class II enhancers appears to have almost identical effect on recall regardless of the method, and indeed the ROC curves are nearly overlapping (**Fig. 1A**). This is particularly noteworthy as the underlying assumption of both models—primarily, feature additivity and independence—are unlikely to be present in the data, yet both perform as well as Random Forests, which does not require such assumptions. Precision-Recall curves also show logistic regression performance closely matches that of random forests, though naïve Bayes precision is poor **(SI Fig 1).** In all cases accounting for heterogeneity increases precision significantly. When the non-enhancer set is purged of enhancers active only post stage 6, the PR curve for RF has AUC > 0.95, demonstrating extremely high sensitivity in the data.

**Figure 1:**
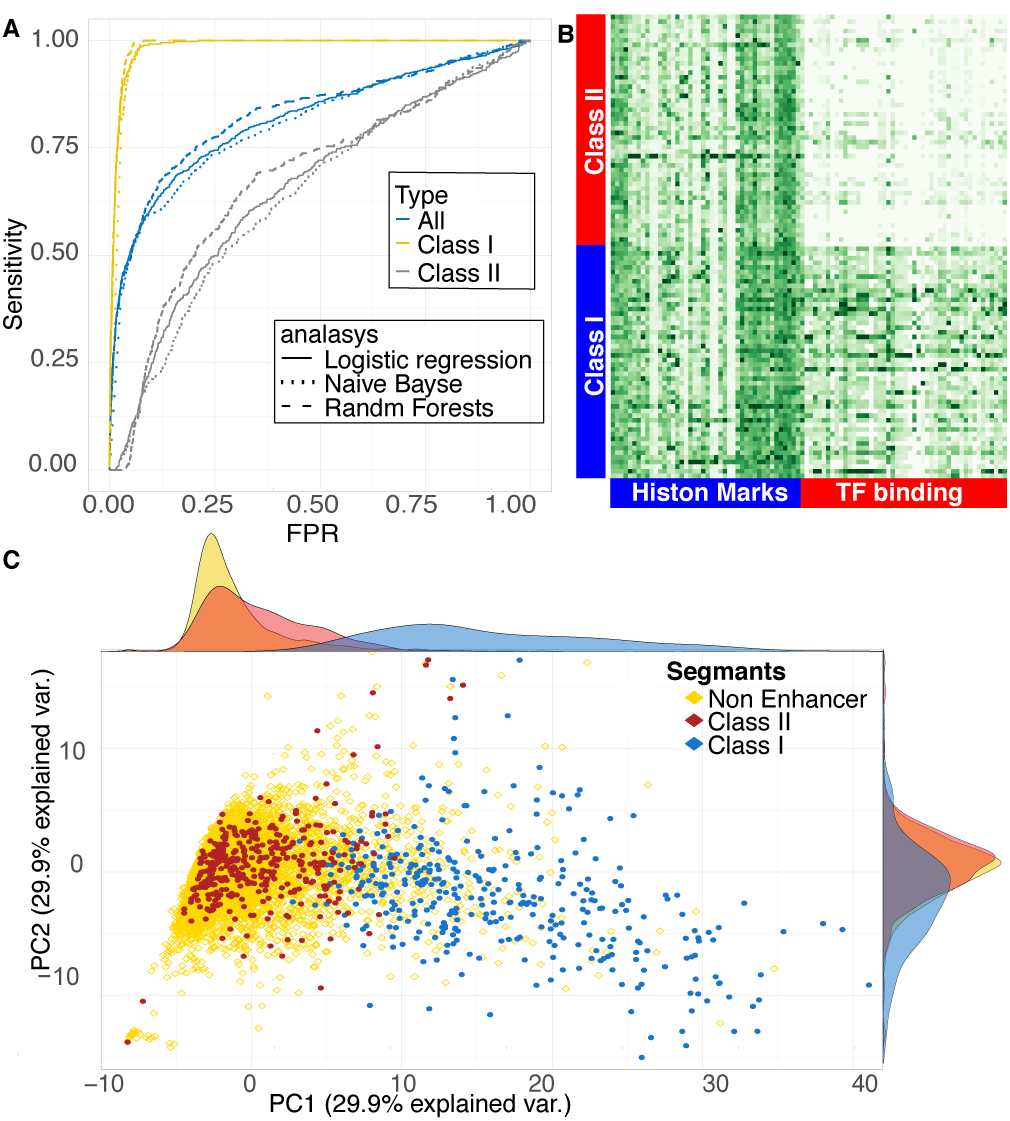
**(A)** Random Forest ROC curves for the complete data set of 7,987 previously validated genomic regions (blue) shows mediocre performance, with an area under the curve (AUC) of 0.83. When only class I enhancers and non-enhancers are used for training, the predictive power rises sharply, AUC 0.99 (yellow). When only class II enhancers and non-enhancers are used, the result is close to a random guess (grey). When predicting the class I enhancer set the ROC curves for Random Forests, logistic regression and a Naïve Bayes classifier are nearly overlapping. **(B)** This can be explained by the co-localization of class II enhancers and non-enhancers in a PCA projection. **(C)** The separation is mainly driven by transcription factors as exemplified by the normalized ChiP strength across features of 200 randomly selected class I and class II enhancers.

This seperation by the model can be understood by Principal Component Analysis (PCA) (**Fig. 1C**): Class II enhancers are collocated with non-enhancers while class I enhancers are separated from both. Examination of feature space statistics of the three groups shows that Class II enhancers are indistinguishable from non-enhancers along our entire feature space— TF DNA binding, histone marks, conservation and DNase accessibility—while class I enhancers segregated from both by multiple features. The separation is most notable in TF DNA binding and DNase accessibility profiles (**Fig. 1B, SI Fig. 2**), where class I enhancers consistently have higher ChIP scores and are more accessible in whole embryo average data. This indicates a possible reason and mechanism for the separation of the two classes and shows that Random Forests can be readily used to separate heterogeneous enhancer sets.

Excluding Class II enhancers from the sampled training set gives us unprecedented prediction accuracy. On a balanced held out, test set, built from genome regions that prior studies suggested half were enhancers and half were non-functional, more than 98% of Class I enhancers are discovered by our algorithm with better than 95% precision. This model would have much lower accuracy if used to predict enhancers genome-wide, however. As one moves away from a balanced test set by adding a more realistic number of inactive genomic regions, the false positive rate in the test set will increase.

To demonstrate this point, Random Forests were trained on a balanced set and then tested on a series of increasingly imbalanced test sets, at various degrees of stringency (**Fig. 2A**). The false positive rate for test sets increases sharply as either the fraction of non-enhancers in the test set increases or as the accuracy of the model—defined during training— increases. This can also be seen in two dimensional plots of the same analysis (**Fig. 2B-C**): unless the sample is very close to a 50%/50% balance, the rise in the false discovery rate in the test set is extremely sharp. Conversely, in genomic scans where non-enhancer regions are at least a hundred fold more prevalent, a precision considerably better then 95% during training (measured out of bag, see Methods) is needed to achieve a 75% false discovery rate in the test.

**Figure 2:**
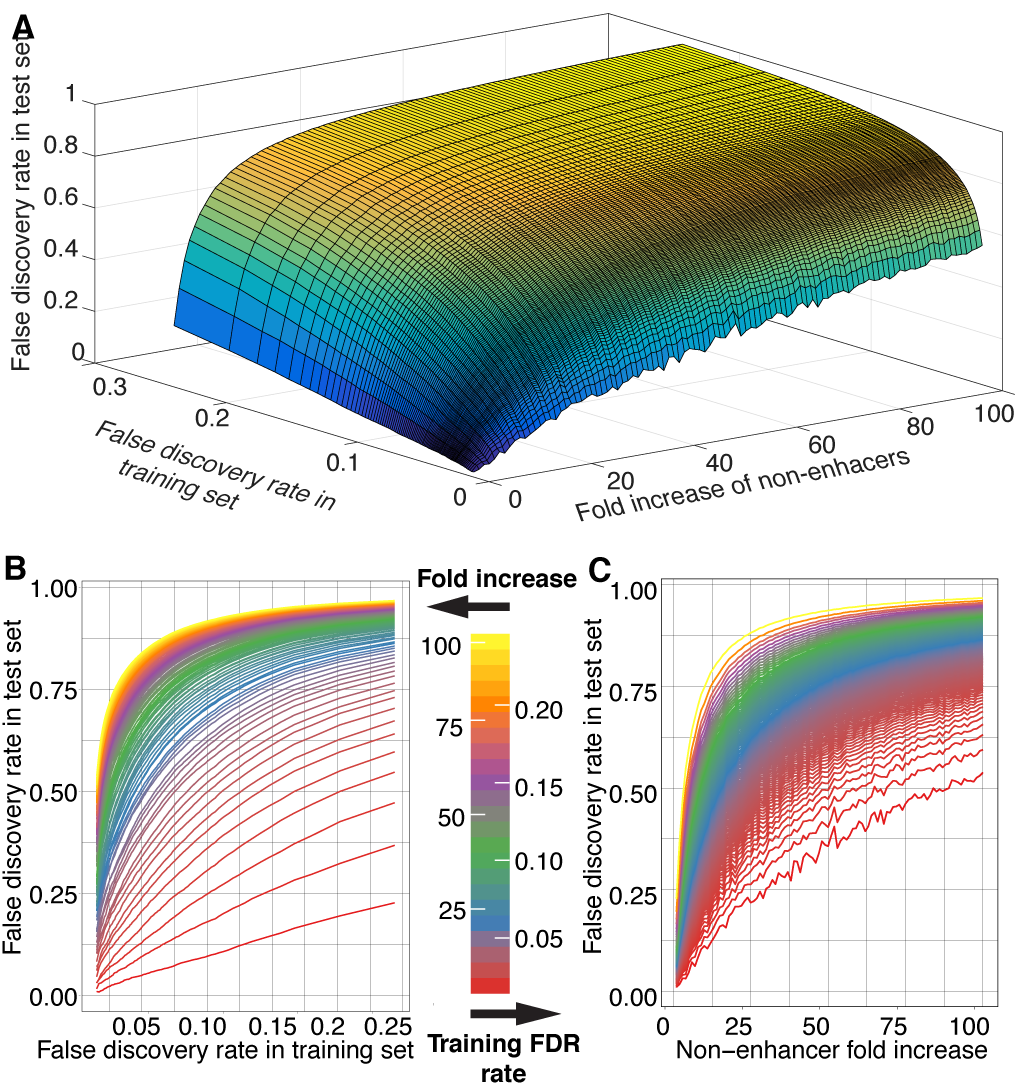
False positive rate is a function of method accuracy and imbalance in the test data. **(A)** A 3-dimensional surface plot shows a sharp increase in the test set false positive rate as either the training set false positive rate or the fraction of non-enhancer regions in the test set increase. This shows that in genomic settings, where the imbalance cannot be controlled, a very high degree of accuracy is required. **(B-C)** Two dimentional plots of the marginals of the 3D image in A, demonstrating the sharp rise in test inaccuracy for both false positive rate in the training set or dilution of enhancer class in the test set.

In the dataset of Kvon *et al.* there are 20 times more annotated non-enhancers than enhancer elements. In randomly drawn test sets with only 5% true enhancers, we find that our fitted model recovers 90% of enhancers with 60% precision. However, our prediction accuracy is likely considerably higher than this analysis implies due to an abundance of false negatives in the high-throughput Kvon *et al.* annotations. Manual reexamination of their reporter gene expression image data for the 100 genomic regions that our method most highly predicted to be enhancers, but which were reported as non-enhancers, revealed that only 15 were true non-enhancers, 47 were clearly enhancers, and the remainder could not be classified due to insufficient data – specifically the lack of embryos of the appropriate stage in the high-throughput images (**Fig. 3C**).

**Figure 3:**
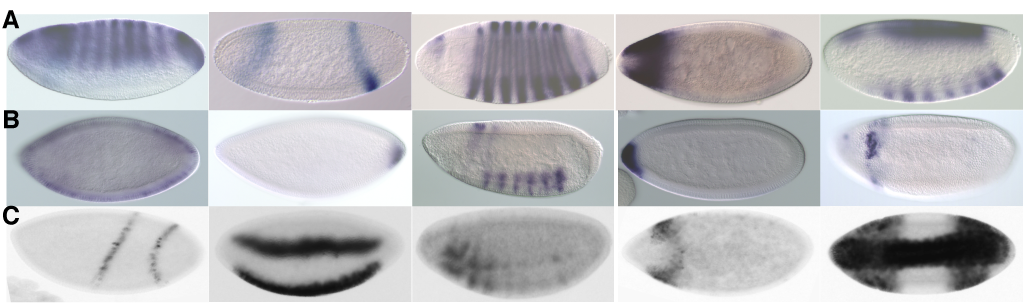
Examples of reporter gene expression patterns driven by **(A)** Class I enhancers, **(B)** Class II enhancers and **(C)** genome regions misclassified by Kvon et al. as non-enhancers in stages 4-6.

### Genome-wide analysis to identify segmentation-driving enhancers in the early embryo

Given the high accuracy of the model on our training and held-out data set, a genome wide search for class I enhancers was feasible. Random Forest was therefore used to predict enhancer probability on a computationally segmented genome (see Methods). More than 82% of all segments had less than 0.01 probability of being enhancer, and more then 93% had less than 0.1 probability **(Fig. 4A**). While it is challenging to see initially as the histogram is dominated by a peak between probabilities 0-0.01, the histogram is in fact bimodal (**Fig. 4A, inset**), with a secondary peak around p = 0.95. To call enhancers a threshold of *p*> 0.75 was established that covers ∼1.6% of the genome and rediscovers *de novo* 98% of the training set. 1,640 Class I enhancers are predicted, 1174 of which do not overlap with training data; 364 overlap known CRM’s identified in a database of enhancers discovered in other studies, Redfly (50-52); and 916 are completely novel. The prediction list can be viewed at (53)

**Figure 4:**
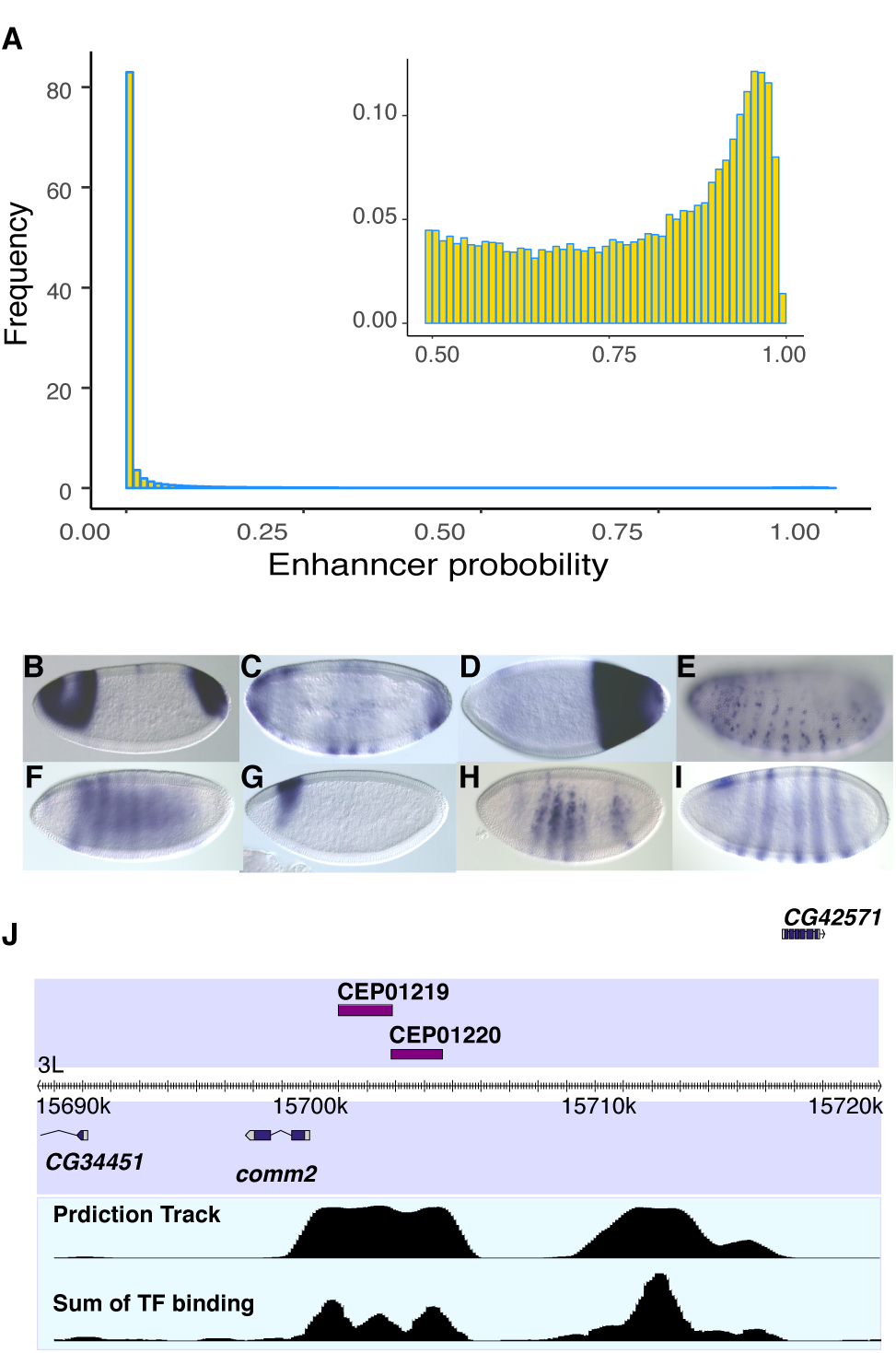
**(A)** Histogram of Random Forest predicted enhancer probabilities for the entire genome. While >82% of the genome has *p* <0.01, a secondary peak can be seen at *p* ∼0.95 (see inset). **(B-F)** As validation, predicted enhancers were inserted into the *Drosophila* genome, and were found to drive spatial expression. **(G,H)** two enhancers are predicted proximal to the *Comm2* gene. Each of their patterns is a component of the *Comm2* expression pattern **(I). (J)** The genomic region of the two predicted enhancers is shown, along with the raw prediction track showing the predicted probability of enhancer activity with 100bp resolution and the sum of TF binding ChIP scores at the same resolution.

To validate the prediction precision, an *in-vivo* reporter expression-driving test was conducted. 5, 11 and 17 genomic regions were selected with probability scores corresponding to estimated (cross validation) local false discovery rates of 4%, 25% and 50% respectively. These FDR were calculated locally in a small probability bound to be tested, rather than for the whole set. Test regions were cloned into the pBPGUW expression vector then injected into flies using the attP integration system (54) (**SI table 3)**. All but three of the enhancers, including all but two of those predicted to be in the 50% FDR region were found to be enhancers (**Fig. 4B-F**). We thus needed to adjust our FDR estimation; assuming a Poisson distribution we obtain a maximum likelihood estimate (MLE) of 12.5% FDR at our previously cross-validation-based 50% FDR threshold (see **methods**). For our entire collection of 1,640 predicted Class I enhancers, we estimate an overall FDR of 13.58%.

An interesting example and validation for the use of transcription factors to separate proximal enhancers (**methods)** can be seen in two predicted segments proximal to the Commissureless gene (*Comm2*) (**Fig. 4J)**, an important protein in axons guidance across embryo’s midline (55, 56). The two predicted enhancers combined expression pattern (**4G,H)** matches the more complicated expression pattern of *Comm2* (**4J).**

### Segmentation driving enhancers (SDE)

We next sought to understand if the separation of the enhancers into two classes in our feature space is related to their biology. In a detailed quantitation of images of embryonic reporter gene expression patterns for 85 randomly selected class I and 82 randomly selected class II enhancers, class II enhancers tend to be expressed in a smaller percent of cells. 74% of class II enhancers are expressed in ≤15% of cells vs only 33% of class I (SI Fig. 3). While separation by this criterion is not complete, it is unlikely that these differences in expression are due to chance (*p*-value < 10^-7^). In addition, we find that class I enhancers are more likely to remain active throughout embryogenesis and show a significant enrichment for the expression in A-P stripes, posterior or gap gene like patterns (*p*-value < 10^-4^). GO-term analysis of the genes proximal to class I enhancers also showed a highly significant enrichment of terms related to segmentation (**Fig. 5**), while those of class II enhancers showed much lower enrichment for any GO terms and no significant enrichment for any particular pathway (**Fig. 5**). We therefore hypothesize that class I enhancers are likely to drive expression patterns needed for establishing the segmented body plan. We thus term class I and class II enhancers Segmentation Driving Enhancers (SDEs) and non-SDEs respectively. We note that while the differences between these two classes are significant, there is not a clear separation in function as a minority of non-SDEs direct patterns of expression resembling those of SDEs (**Fig. 3A,3B**).

**Figure 5:**
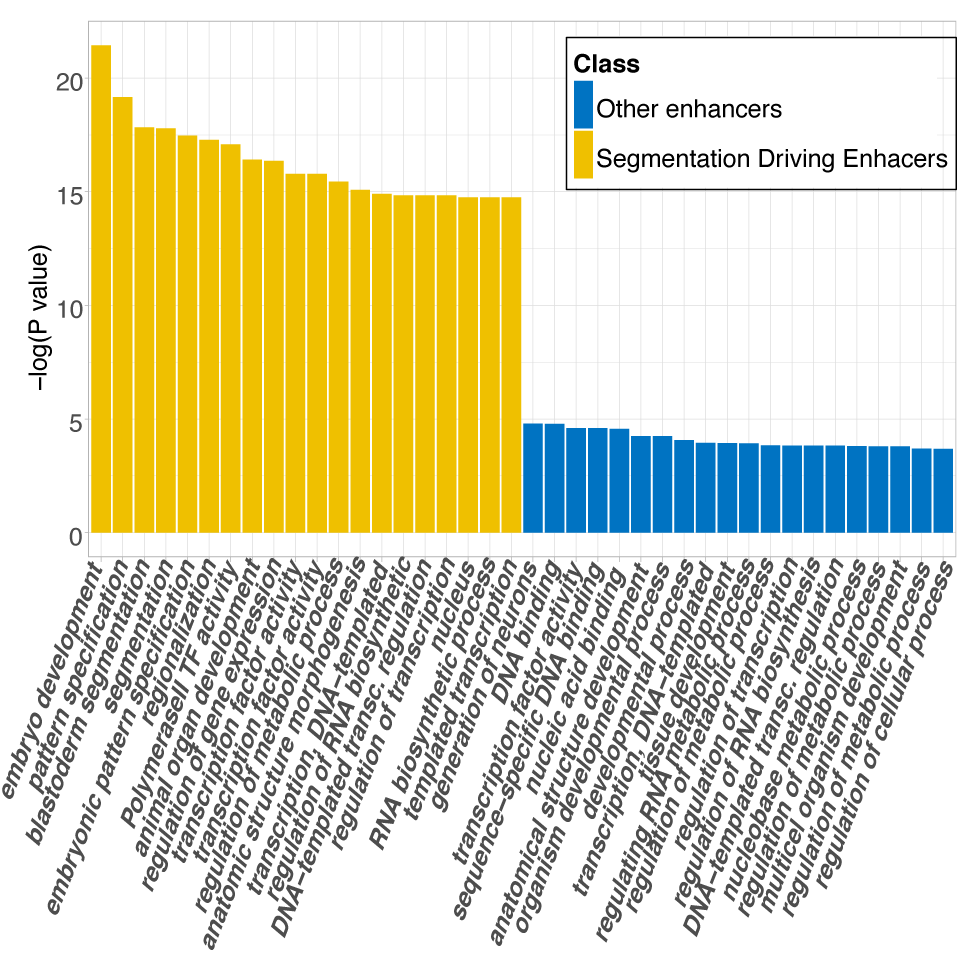
The significance (measured as the negative log of the *p*-value) of GO-term enrichment in genes proximal to class I enhancers is very high in terms associated with development and segmentation (SDES, yellow). For Class II enhancers no significant GO-term enrichment (P-value below 10^-5^) is found (non-SDEs, blue). PLEASE CHANGE THE LABEL ON THE FIGURES TO SDEs AND NON-SDEs TO BE CONSISTENT WITH THE REST OF THE TEXT.

### Feature importance is dominated by transcription factors

The Random Forest importance measures “mean decrease accuracy” and “mean decrease Gini” (49) varied between bootstrap repetitions, but in all cases a small set of TFs were found near the top of the importance ranking list. This can be seen by the spread of the bootstrap confidence interval of these two importance measures calculated in 50,000 trees (**Fig. 6A, SI Fig. 4A**). The sum of TFs binding per genomic region, along with individual the ChIP binding scores for several TFs (Kr, Med, Twi, Dl, D) were the most important by both measures, and were also the most often used by the model **(SI Fig. 4B**). Other TFs, such as Bcd and Ftz (33), were uninformative despite their importance for embryo segmentation. This can be at least partially explained by low coverage in the ChIP-chip data, as there is a clear correlation (r = 0.7) between coverage and importance measure (**SI Fig. 5a**). The only histone mark to have an importance above random noise was H3k4 mono-methylation (H3K4me1). All 40 other histone and histone modifications, including the H3K27 acetylation (H3K27ac) that has been widely regarded as a key indicator of enhancer regions **(18, 57)** were found uninformative by the model in the presence of the transcription factor data.

**Figure 6:**
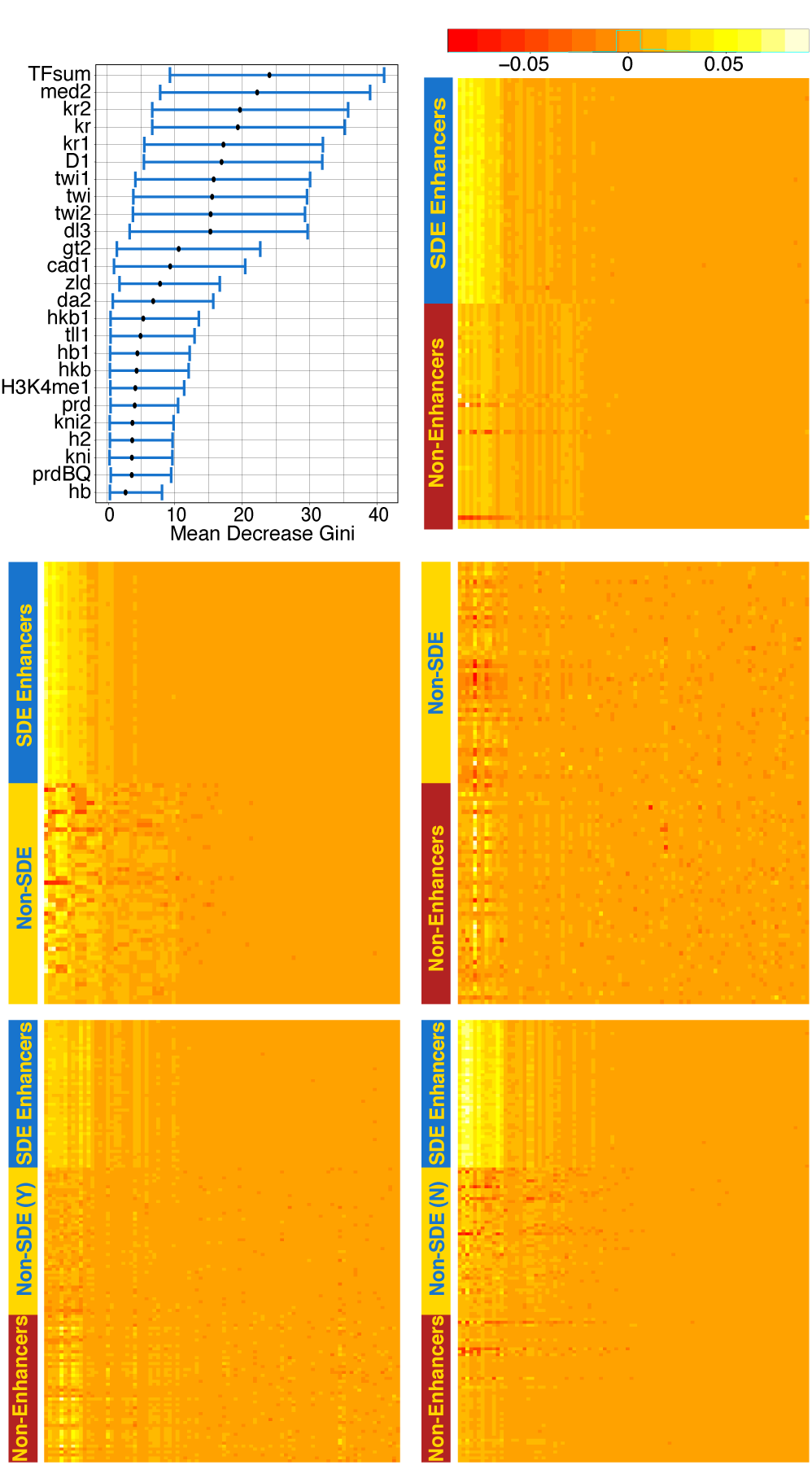
**(A)** Feature importance is dominated with transcription factors, with the H3K4me1 the only histone mark in the top 25. **(B-F)** “Local importance” measurements of randomly selected segments indicting how important each feature was in the segment classification when forest was trained on **(B)** SDEs vs. non-enhancers; **(C)** SDE vs. non-SDEs; **(D)** Non-SDEs vs. non-enhancers; **(E)** SDEs and non-SDEs vs. Non-enhancers; **(F)** SDEs vs. non-SDEs and non-enhancers. Feature order (x-axis) can be found in **Supplementary text**

### Localized feature-importance measures support a two-class structure

Random Forest’s “local importance” is a third measure that provides a detailed determination of the importance of each feature in classifying each instance, allowing a more direct understanding on the Random Forests decision making process (**Fig. 6B-F).** This measure shows that the same small set of fetures are used to distinguish SDE and non-enhancers (**Fig. 6B**) as are used to distinguish SDEs from non-SDEs (**Fig. 6C**), while an attempt to seperate non-SDEs and non-enhancers (**Fig. 6D**) shows that no variable can consistantly be used and that many more parameters are employed. The increase in featues used and the blurring of decision-criteria is also seen when non-SDEs are presented to random forest as enhancers (**Fig. 6E**) rather than as non-enhancers (**Fig. 6F**). Spectral clustering is a technique that relies on the eigenvector of the similarity or afinity matrix projection of the data, usually followed by k-nearest neighbors or k-means clustering (58). It is an efficiant way of dimension reduction, and the number of clusters in the data can often be infered by the eigenvalues. Applying spectral clustering to our data fails to separate SDE from non-SDEs, and the eigenvalues of the affinity matrix indicates a single cluster. Applying spectral clustering to the local importance matrix yields a good separation of the data, however, with a sharp jump after the second eigenvalue (**SI Fig. 5b**) – consistent with the presence of a two-class structure.

## Discussion

The identification of enhancer elements from genomics data has remained a challenging problem—in part due to the relative scarcity of enhancers in genome sequences vs non-enhancer sequences. As we illustrated in Figure 2, even an incisive enhancer prediction algorithm fitted on balanced training data (i.e. a training set with nearly equal numbers of positive and negative elements) is likely to generate high false discovery rates when tested on a genome-wide scan. Hence, to accurately discover enhancer elements using *in silico* techniques, extremely high fidelity models are needed.

Though high precision predictions were reported previously, the validation methods and measures used in the literature varied greatly. Many papers defined success as the colocation of data for epigenetic marks such as p300 and H3K27ac, but it is yet to be established that these marks are exclusive to enhancers or that all enhancers possess them. Indeed, we report here a class of H3K27ac-free enhancer (SI Table 4). Other reports tested for functional enhancer activity of genomics regions from the top of a rank list (17), which gives a biased estimate of the overall prediction accuracy. We suggest that precision must be measured by testing throughout the prediction rank list to establish a uniform unbiased measure of success.

We found that the prediction of enhancer elements *en masse* was vexed by heterogeneity among enhancer elements. For about half of previously validated enhancer elements, strong TF *in vivo* DNA binding signals for multiple factors is indicative of enhancer activity. The remaining half of validated elements are typically bound more weakly by fewer TFs (Figs. 1B and S1). For this latter set, the residual TF binding signal is only weakly associated with enhancer activity. That is, a prediction engine that works extremely well on one class tends to fail on the other. We posit that this challenge—heterogeneity in element classes—is a widespread and foundational challenge in genomics. For example, the emerging literature on “chromatin priming elements” (59) demonstrates the existence of “enhancer-like” functional elements that, while they share chromatin structure and similar patterns of transcription factor occupancies with enhancers, do not themselves drive patterned expression—rather they establish chromatin context that subsequently gives rise to enhancer activity for proximal elements. It may be that the class of elements we presently denote “enhancers” is in fact diverse, admitting elements that exert regulatory effects through a variety of underlying molecular mechanisms. Indeed, it remains unclear what fraction of enhancers require eRNAs for their activity (59), or whether priming elements are transcribed like many enhancers.

It may also be that the non-SDEs or “class II” enhancers we study here are simply regulated by cohorts of TFs we have yet to survey. These enhancers, however, are often expressed in a smaller fraction of the embryo than SDEs and have lower whole embryo average DNase I-seq and TF ChIP biding scores (SI Fig. 1), consistent with them being accessible in only a small subset cells. This would thus make non-SDEs less amenable to interrogation through whole-animal ChIP-seq. The differences between the two enhancer classes are statistical, though, not categorical. While there is a significant enrichment in segmentation GO-terms in SDEs compared to non-SDEs, some non-SDEs also display segmentally repeating expression patterns similar to those of SDEs, and many (20%) showing activity above the 15% expression area threshold, and 8.5% were active in as much or more of the embryo as the median for SDEs.

Our validation assays revealed that cross-validation had led to significant overestimation of the false discovery rate for SDEs. We attribute this to an abundance of false negatives in our training set as our analysis indicated that ∼10% of negatives are erroneously labeled, which would double the number of positives and explain the significant increase in validated false discovery rates we observed. An alternative explanation for the discrepancy is selection bias in our training set as the genomic regions tested in the previous studies whose data we used were not selected at random. Thus, it is possible that there is a clearer separation of features when the full selection of enhancers in the whole genome are considered. Overall, we recover 98% of the training set SDEs with an estimated false discovery rate of less than 15%, indicating that our genome-wide predicted catalog of these elements may be close to comprehensive. Further experiments, particularly concentrated at high false discovery rates, are needed to better assess the boundary between functional and non-functional elements. At this time, it appears that at least 1600 elements, composing more than 1.6% of the *Drosophila* genome, are involved in establishing early body patterning in the blastoderm.

## Material and methods

### Data acquisition and processing

25% FDR Transcription factor ChIP-chip data was taken from the *Drosophila* TF network project (available on http://genome.ucsc.edu/cgibin/hgTracks), containing data for 22 transcription factors: BCD, CAD, D, DA, DL, FTZ, GT, H, HB, HKB, KNI, KR, MAD, MED, PRD, RUN, SHN, SLP, SNA, TLL, TWI, Z, some with biological duplicates to give 34 tracks. 1% FDR ChiP-chip data for polII binding was also taken from BDTNP (37-39).Histone, histone modification and zld chip-seq data were retrieved from UCSC genome browser track provided by Li el al.(40). Histone modifications data collected in Zld mutant strain were not used, all other tracks were included in the analysis. DNase accessibility data and 12-fly conservation phastCons scores were obtained from UCSC genome browser(60-62), as was FlyBase gene data for exon, coding exons and intron location data(63). Bidirectional RNA transcript data was obtained from (64), and analyzed as described in (65). Transcription start site was taken as the start of the first exon in Flybase’s mRNA data described above. All data and analysis was done using R5 dm3 genome annotations.

Though 80% of the DNA segments in the training set were between 2-2.5Kb long, segment sizes varied from 100bp to 4.5KB in the set, and the percent of enhancer region contained by each segment is unknown, making averages a biased estimator. Thus, the maximum of ChiP data was calculated over every segment in the training set and the segmented genome using bedtools and UCSC genome browser utilities for TF data, histones, conservation score and DNase accessibility. In addition, the sum of TF biological replicas and the sum of all TF tracks was also calculated and included as features in the model. In addition to the maximum score, for Zld higher-resolution ChiP-seq data and for the conservation phastCons conservation scores we also calculated the average over the segment, maximal score over a sliding window of 200,500 and 1000bp, and the longest continues stretch of scores above the 0.85 quantile. For the gene data, bedtools coverage was used to calculate percent of segment covered by exons, coding exons or introns andand 3 binary tracks indicate the presence or absence of intron and exons. Bedtools closest to calculate distance to the closest tss and to polII binding peak.

### Modeling

Random Forests were modeled in R (66) using RandomForest(48). Initial feature set culling was done through error rates average of 1000 forests of 500 trees when excluding/adding one feature at a time. Our training data is highly unbalanced, with only 10% of segments being enhancers. To improve Random Forest performance balanced samples were used as training set. To improve stability of the prediction, and counteract the sampling process employed by balancing the training set, we relied on forest voting. 1000 Forest of 50 trees each were trained on a randomly selected sets of 300 enhancer and 300 non-enhancers with 10% of the data held out of the samples and used as test set. The fraction of trees in all forests voting for each segment serving as probability of being an enhancer. this was repeated until such score was computed for each segment in the set. The same sampling and testing scheme was employed for logistic regression, and naïve Bayes (67).

Importance measures varied from sample to sample and averages required 10,000 Forest of 50 trees to converge. To increase stability of the importance measure, the average of 50,000 Random Forests mean decrease in accuracy and mean decrease in Gini index were used to find the importance Random forests confidence intervals. For local importance calculations we used a single forest of 50,000 trees produced using all enhancers and a balanced non-enhancers subsample.

### Analyses

ROC curve areas were calculated with R package PRROC (68). PCA was done using prcomp (66). Go term analysis used bedtools (69) to find FlyBase genes located inside training enhancer regions, or to identify the closest genes if none are overlapping. David bioinformatics resource (70, 71) was used to find and quantify Go term and Go-term enrichment, with the full set of ∼8000 genomic regions as the genomic background. To find Affinity matrix of the data we converted Euclidian distance into a similarity matrix, and calculated 7 nearest neighbors for each segment. Spectral clustering and eigenvalue extraction was done using kknn (72) with default settings. We used a masked strategy to assess expression size and pattern on an unannotated randomly ordered set of both enhancer classes.

### Genome wide prediction

A sliding window of 1000bp with 100bp distance was used to create segments of the entire drosophila genome, and 1000 trained on SDE and non-enhancers only with our usual sampling scheme was used to predict enhancers genome wide, with the %trees taken as probability. For each 100bp segment the average of the overlapping segments was calculated, and those above the 0.75 threshold were kept. adjacent segments were merged. Segments longer then 1.5Kb were separated based on peaks in the sum of transcription factors data when possible: The normalized sum of transcription factor binding was calculated for each 100bp window, second derivative used to detect peaks, and peaks closer then 200bp merged. If more than one peak remained, the minima between adjacent peaks was used to separate the longer predicted enhancers. Once boundaries were established, the genomic prediction scheme was repeated to establish probability of the entire enhancer. New false discovery rate MLE and confidence intervals were calculated under an assumed Poisson distribution, one per predicted FDR range. A second order polynomial was fitted to these 3 points together with a forth of 100% FDR at 0 probability as the threshold. The 1640 predicted enhancers were fitted to the polynomial, with the average taken as the predicted and CI FDR score.

### PCR of Fragments from Genomic DNA and Cloning into the Gateway Vector

*PfuUltra* High-Fidelity DNA Polymerase (Stratagene) or EASYA DNA polymerase (Agilent) was used to amplify selected fragments (see above) by using isogenic genomic DNA from *y*; *cn bw sp* (Adams MD *et al.* (2000) The genome sequence of *Drosophila melanogaster*. *Science* 287:2185–2195) as a template. The PCR products were confirmed by agarose gel analysis, purified by using the QIA-quick PCR Purification Kit (Qiagen). PCR fragment cloning was performed by adding 3 A overhangs to the PCR products produced using the PfuUltra High-Fidelity DNA Polymerase (A overhangs were not added to the products produced using the EasyA DNA Polymerase) with the addition of dATP and *Taq* polymerase in a 10-min incubation at 72°C before Qiagen purification. The products (9.5 ul of each) were used in a TA TOPO cloning reaction with pCR8/GW/TOPO as described by the manufacturer (Invitrogen). Cloning reactions were allowed to proceed for 30 min at room temperature, and then 2 ul of each reaction was used to transform Mach1 cells (Invitrogen). For each cloning reaction, two isolates were picked, purified, and confirmed by sequence verification.

### Sequence Verification of Clones

Two Gateway clones were picked for each enhancer fragment, for a total of 78 processed clones. Sequencing primers M13 FWD −20 (Invitrogen; 5’ GTAAAACGACGGCCA 3’) and M13 REV (Invitrogen 5’ GGAAACAGCTATGACCATG 3’) were used to generate sequence to verify targets. One clone was identified and selected for future studies.

### Transfer of Gateway Clones into Integration Vectors

Thirty-seven nanograms (ng) of the destination vector, pBPGUw, were combined with 37.5 ng of DNA carrying a PCR fragment cloned in the Gateway vector in a LR reaction (Invitrogen) and incubated overnight at room temperature. TAM1 cells (Invitrogen) were transformed with 2.5 ul of the LR reaction and plated. A single isolate from each reaction was picked into a 96-well Beckman Deepwell block, allowed to grow overnight at 37°C and DNA was prepared by using the PerfectPrep kit (5 PRIME). The constructs were verified by analysis of restriction enzyme digests. A second isolate was picked in cases where there was a discrepancy between the observed and expected results. DNA for injection was prepared from 7 ml of overnight culture for production of transgenic flies.

### Drosophila Genetics

DNA constructs (100 –200 ng/ ul) were microinjected into embryos derived from parents homozygous for both the *attP2* integration site (Groth AC, Fish M, Nusse R, Calos MP (2004) Construction of transgenic *Drosophila* by using the site-specific integrase from phage phiC31. *Genetics* 166:1775–1782.) and a fusion gene encoding the *PhiC31* integrase under the control of the *nanos* promoter (*nos–integ*), which provides a maternal source of integrase (Bischof J, Maeda RK, Hediger M, Karch F, Basler K (2007) An optimized transgenesis system for *Drosophila* using germ-line-specific phiC31 integrases. *Proc Natl Acad Sci USA* 104:3312–3317.). Single males derived from these embryos were crossed to *y w;Sco/CyO* females, and males carrying the inserted construct (identified by their *w+* eye color) were selected; integrase is removed in this step. These males were crossed to *w[1118]/Dp(1;Y)y[+]; TM2/TM6C, Sb[1] (Bl stock # 5906)* females to establish balanced, homozygous stocks. We obtained 48 transgenic lines containing predicted enhancer elements called CEPs.

### Verification of Insertion Site and Fragment Identity by Genomic PCR

To verify the identity of transformant flies and to confirm that all integration events occurred at the *attP2* site, we performed genomic PCR on DNA isolated from homozygous transformant f lies. Twenty f lies were homogenized and genomic DNA isolated by using the ZR Genomic DNA II Kit (Zymo Research). To assay proper integration in the *attP2* landing site, PCR was performed by using a primer from the *y* gene marker in the *attP2* genomic docking site (TCATGACTTTGTTGCCTTAGA) and a reverse primer from the *w* gene (CGAAAGAGACGGC-GATATT) carried in the constructs. Only proper integration events would yield a product of 1839 bp, because *y* and *w* lie more than 2 Mb away in the *Drosophila* genome. Fragment identity was confirmed by using a vector-specific primer (ACAAGTTTG‐ TACAAAAAAGCAGGCT) and a reverse primer specific to the cloned fragment being tested for enhancer activity; the position of the fragment-specific primer was chosen so as to yield a PCR product of 350 to 400 bp.

### Embryo Whole Mount in Situ Hybridizations

Embryos were collected directly from the homozygous stock. Embryonic whole mount *in situ* RNA hybridizations were performed as described previously (PMID:19360017). A summary is shown in Supplemental Fig. 6.

## Acknowledgements

This work was funded by NIH/NIGMS grant 5P01GM0999655 under Department of Energy contract no. DE-AC02-05CH11231. JBB’s work was supported by NHGRI R00 HG006698 and Department of Energy/LDRD DE-AC02-05CH11231/14-200. PJB was supported by 1U01HG007031-01. SEC was also supported by NIH grant NIH R01-GM076655.

## Author Contributions

JBB, MDB and SEC designed the project. JBB managed computational analysis. SEC managed data collection. HA developed, conducted, and interpreted analyses. HA designed and generated figures. HA oversaw data organization and homogenization across platforms. KHW designed the primers, RW and SP performed the PCR, generated and validated clones and made DNA for injections, SP and BF made transgenic animals and homozygous stocks, SK prepared fixed embryos, RW performed hybridizations and RW, BF and AH collected and analyzed the image data. HA and OSS analyzed data. HA and JBB wrote the manuscript with input from all authors, and SEC and MDB contributed methods text. All authors discussed the results and commented on the manuscript.

